# Phylogeographic analysis of shrubby beardtongues reveals range expansions during the Last Glacial Maximum and implicates the Klamath Mountains as a hotspot for hybridization

**DOI:** 10.1101/2020.10.06.328757

**Authors:** Benjamin W. Stone, Andrea D. Wolfe

## Abstract

Quaternary glacial cycles often altered species’ geographic distributions, which in turn altered the geographic structure of a species’ genetic diversity. In many cases, glacial expansion forced species in temperate climates to contract their ranges and reside in small pockets of suitable habitat (refugia), where they were likely to interact closely with other species, setting the stage for potential gene exchange. These introgression events, in turn, would have degraded species boundaries, making the inference of phylogenetic relationships challenging. Using high-throughput sequence data, we employ a combination of species distribution models, models of demographic history, and hybridization tests to assess the effect of glaciation on the geographic distributions, phylogenetic relationships, and patterns of gene flow of five species of *Penstemon* subgenus *Dasanthera*, long-lived shrubby angiosperms distributed throughout the Pacific Northwest of North America. Surprisingly, we find that rather than reducing their ranges to small refugia, most *Penstemon* subgenus *Dasanthera* species experienced increases in suitable habitat during the Last Glacial Maximum. We also find substantial evidence for gene exchange between species, with the bulk of introgression events occurring in or near the Klamath Mountains of southwestern Oregon and northwestern California. Subsequently, our phylogenetic inference reveals blurred taxonomic boundaries in the Klamath Mountains, where introgression is most prevalent. Our results question the classical paradigm of temperate species’ responses to glaciation, and highlight the importance of contextualizing phylogenetic inference with the demographic histories of the species of interest.

## Introduction

Throughout the Quaternary Period, the Pacific Northwest of North America (PNW) experienced dramatic shifts in climate due to the repeated expansion and retreat of glaciers (Shafer, Cullingham, Côte, & Coltman, 2010). At the peak of the most recent glacial expanse, known as the last glacial maximum (LGM), global temperatures in the PNW were substantially colder than present-day (Otto-Bliesner & Brady, 2006), and massive ice sheets rendered large regions of land inhospitable for many species (Pielou, 2008). After the LGM, conditions in the PNW became increasingly hot, reaching a temperature maximum near the mid-Holocene period, about 6,000 years BP (Renssen *et al*., 2009; Wanner *et al*., 2008).

Such dramatic fluctuations in climate undoubtedly altered species’ distributions through time. Many species may have survived in one or more pockets of suitable habitat (i.e. refugia) during these glacial cycles. Given the myriad effects that such processes could have on the spatial distribution of contemporary genetic diversity, many studies have focused on identifying such refugia and better understanding patterns of contraction and expansion in response to glacial cycles (Avise, 2000; Hewitt, 2000). As a result, there is a rich history of phylogeographic research in the PNW (Shafer *et al*., 2010). Although the response of species to glacial cycles depends on their ecological and climatic tolerances (Hewitt, 2004; Stewart, Lister, Barnes, & Dalén, 2010), phylogeographic studies have identified several recurrent patterns of genetic differentiation across a broad range of taxa. For example, Soltis, Gitzendanner, Strenge, and Soltis (1997) described a north-south pattern of genetic differentiation in several species of plants distributed along the Cascades and Coastal Mountain ranges. In addition to this, Soltis *et al*. (1997) identified reduced genetic diversity in the northern portion of some species’ ranges compared to the south, suggesting the presence of southern refugia for many species during the LGM, and the potential for multiple refugia in the coastal range and Klamath Mountains of southwestern Oregon and northwestern California. Brunsfeld, Sullivan, Soltis, and Soltis (2001) elaborated on these findings, outlining expectations for the hypotheses outlined in Soltis *et al*. (1997), and formulating new phylogeographic hypotheses for species with Cascade/Sierran distributions (*e.g*., the clinal environment hypothesis).

The Klamath Mountains, one of the potential refugial locations highlighted by Soltis *et al*. (1997), host a complex vegetative history owing to their old geologic age, edaphic diversity, and their ability to support mesophytic and xerophytic plant communities (Whittaker, 1961). The Klamath region has long been considered a potential refugium for plant species, as species with more northernly distributions likely invaded the Klamath Mountains during the cooler conditions of the Pleistocene, then remained there once the climate warmed again by moving to higher elevations (Smith & Sawyer, 1988; Whittaker, 1961). Indeed, the Klamath Mountains have been identified as an important geographic feature for many plant taxa, including as an area with genetic substructure or divergence (Furnier & Adams, 1986; Soltis *et al*., 1997), as a major phylogeographic break point or area where sister species’ ranges abut (Gugger, Sugita, & Cavender-Bares, 2010; Patterson & Givnish, 2003), and as a glacial refugium (Eckert, Tearse, & Hall, 2008; Kiefer, Dobeš, Sharbel, & Koch, 2009). The biogeographic importance of this region is not just limited to plants, however, as it is also a potential refugium for *Plethodon* salamanders (Pelletier, Duffield, & DeGrauw, 2011), *Anaxyrus* toads (Goebel, Ranker, Corn, & Olmstead, 2009), and *Taricha* newts (Kuchta & Tan, 2005).

Advancements in sequencing technology have given rise to an increased availability of high-throughput sequence data for use in phylogeographic studies (Garrick *et al*., 2015), which has in turn allowed researchers to compare more complex models of demographic history (*e.g*., Smith *et al*., 2017). Although it has long been suggested that the expansion and contraction of species ranges in response to glacial cycles led to the formation of many hybrid zones (Hewitt, 2011; Stebbins, 1985), the use of high-throughput sequence data in phylogeographic studies has increased awareness of the prevalence and complexity of gene flow and hybridization in species’ responses to climatic change (Maier, Vandergast, Ostoja, Aguilar, & Bohonak, 2019; Ruffley *et al*., 2018; Smith & Carstens, 2020). There are numerous potential outcomes of secondary contact, but some well-known examples include lineage fusion, (Petit *et al*., 2003), adaptive introgression (Anderson & Stebbins, 1954), and speciation via reinforcement (Butlin, 1987). In cases where lineages from separate refugia hybridize upon secondary contact, genetic diversity may increase as the result of lineage fusion (Maier *et al*., 2019; Petit *et al*., 2003), although in some cases there may be a loss of genetic diversity, instead (Colella *et al*., 2018). Hybridization after substantial climatic changes could have adaptive benefits, as novel gene combinations produced upon secondary contact could be beneficial in new, open environments (Stebbins, 1985). The onset of hybridization often does not have adaptive advantages, however. In cases where hybrids are less fit in their environment than either parent, hybridization may facilitate speciation via reinforcement (Butlin, 1987; Dufresnes *et al*. 2020). Hybridization could also be due to the unusual circumstances surrounding the colonization of new habitats. For example, colonizers at the front-end of expansion are more likely to hybridize due to limited mate choice and a multitude of new contact points with closely related species (Currat, Ruedi, Petit, & Excoffier, 2008).

In this study, we aim to understand the response of five species of the genus *Penstemon* Schmidel (Plantaginaceae) to climatic fluctuations during the late Quaternary period, in order to better appreciate how geographic distributions and demographic histories may align to promote gene flow between species. The genus *Penstemon*, commonly known as the beardtongues, is the largest genus of angiosperms endemic to North America, with nearly 300 described species (Freeman, 2019; Wolfe *et al*., 2006). Owing its species richness to a putative adaptive radiation, *Penstemon* exhibits exceptional floral diversity, and its species occupy a wide variety of ecological niches, although in general, they prefer semi-disturbed, arid habitats (Wolfe *et al*., 2006). Although some phylogenetic relationships between *Penstemon* species are obscured, likely due to incomplete lineage sorting (Wessinger, Freeman, Mort, Rausher, & Hileman, 2016; Wessinger, Rausher, & Hileman, 2019), one consistent taxonomic group has been the subgenus *Dasanthera*, which contains nine species total, and is sister to the rest of the genus. Known colloquially as shrubby beardtongues, members of *Penstemon* subg. *Dasanthera* are primarily outcrossing, long-lived plants, typically persisting as low-lying subshrubs in semi-disturbed, rocky habitats. Dispersal is apparently limited – seeds have no obvious mechanisms to facilitate wind-, water-, or animal-mediated dispersal – and this is thought to contribute to their propensity to form scattered, isolated populations (Every, 1977). Species in *Penstemon* subg. *Dasanthera* are found mainly in mountainous areas of the PNW, but extend into surrounding regions, including California, western Montana, northwestern Wyoming, northern Utah, and western Nevada. Four species (*P. rupicola*, *P. cardwellii*, *P. newberryi*, and *P. davidsonii*) have a primarily Cascades/Sierra Nevada distribution, three species (*P. lyallii, P. ellipticus*, and *P. montanus*) have a northern Rocky Mountains distribution, and one species (*P. fruticosus*) is distributed in both the Cascades and northern Rocky Mountains, and in scattered mountains surrounding the Columbia Basin. Hybridization is common in *Penstemon* subg. *Dasanthera*, and there are many well-documented localities at which natural hybrids form, both at local scales, where persistent backcrossing into parental species is unlikely, and at wider scales, wherever species distributions overlap (Clausen, Keck, & Hiesey, 1940; Datwyler & Wolfe, 2004; Every, 1977). Of particular interest in this context are the Klamath Mountains. Every (1977) identified this area as a hotspot for *Penstemon* subg. *Dasanthera* hybridization, noting the overlap of several species’ distributions, and suggesting that hybridization was likely ancient rather than the result of recent and localized introgression. The goals of the present study are to (1) estimate relationships among species of *Penstemon* subg. *Dasanthera* (hereafter referred to as *Dasanthera*) found in the Cascades and Sierra Nevada mountains, (2) identify the location and timing of refugia for these species, and (3) identify introgressed *Dasanthera* lineages at a broad geographic scale, focusing on the Klamath Mountains.

## Materials and Methods

### Data generation

We collected a total of 141 samples representing *Dasanthera* species found in the Cascade and Sierra Nevada Mountains (*P. rupicola, P. cardwellii, P. newberryi, P. davidsonii*, and *P. fruticosus*), and 3 samples of *Penstemon montanus* var. *montanus* from Idaho for use as an outgroup. Because our goals for this study were to better understand the demographic histories of species found on the western side of the Columbia Basin, we did not include samples of *P. lyallii* and *P. ellipticus*, as these species are distributed only in the northern Rocky Mountains, east of the Columbia Basin, and are thus outside the immediate scope of this study. For this reason, we only included samples of the widespread and variable *P. fruticosus* from the Cascades Mountains, rather than including samples from the Rocky Mountains. In addition, we did not include samples of the rare and narrowly endemic *P. barrettiae*, which is a species of conservation concern in the states of Washington and Oregon, and is found only along a roughly fifty mile stretch of the Columbia River east of Portland, OR. The majority of samples (97) were collected during the summers of 2016, 2017, and 2018. The 27 remaining samples collected by the authors were collected either in 1996 or 1999. We also included samples from 20 herbarium tissue loans from herbaria at the University of Washington and Oregon State University. These samples ranged in collection date from 1993 to 2017. In total, our sampling represents 86 unique localities for five of the six *Dasanthera* species and four of the five varieties present in the Cascade and Sierra Nevada Mountains, across the bulk of the range of most of these species (Supplemental Table 1).

All samples collected by the authors were dried with silica gel immediately upon collection. Leaf tissues from herbarium samples were procured directly from herbarium sheets, or from additional pouches of dried material accompanying the collections when available. DNA was extracted using a modified CTAB protocol (Wolfe, 2005) and quantified using a Qubit fluorometer. We prepared Genotyping by Sequencing (GBS) libraries using 100 nanograms of DNA from each sample and a modified version of the Elshire *et al*. (2011) protocol. We sequenced DNA libraries on an Illumina Hi-Seq 2500 using paired-end 150 bp sequencing at Novogene Corporation Inc. (Sacramento, CA).

We used *ipyrad* v0.9.20 (Eaton & Overcast, 2020) for GBS data processing. We trimmed all reads to 50 bp prior to analysis and discarded reverse reads from the paired-end sequencing, because preliminary analysis indicated that doing so produced higher-quality loci with more overlap across species. We produced six different types of data sets with *ipyrad*. The first data set includes every sample in the analysis, including the outgroup. The remaining five data sets are species-specific, *i.e*., each data set only includes samples from a single species. All data sets only include loci that are present in at least 50% of the samples in that data set, and are limited to a maximum depth of 100,000 reads. The remainder of the parameters for data processing are the default parameters for *ipyrad*.

### Identification of populations

We used STRUCTURE (Pritchard, Stephens, & Donnelly, 2000) on the species-specific data sets to estimate the number of populations in each species, and to assign individuals to putative populations. We implemented STRUCTURE with an iterative approach using the *analysis.structure* command in *ipyrad*. For each data set we discarded the first 250,000 generations as burn-in, and ran STRUCTURE for 1,000,000 steps thereafter, performing ten replicates for each value of K tested. All other parameters were left on the default values set by the *ipyrad.analysis* toolkit. We considered values of K=1 to K=10 for each species, except for *P. fruticosus*, for which we considered values of K=1 to K=6 due to its smaller sample size (n = 10). We then used visualizations of log-likelihood scores and the ΔK method (Evanno, Regnaue, & Goudet, 2005) to determine the optimal value of K, and visualized results using the *ipyrad*-analysis toolkit. All downstream analyses that implement the results of STRUCTURE do so by assigning individuals to the highest percentage genetic cluster in the most likely value of K for that species (except for *P. newberryi:* see Results section).

### Species distribution models

To better understand how species’ distributions have changed through time, we built species distribution models (SDMs) for the present-day, the mid-Holocene warm period, and the LGM. To obtain localities to build SDMs, we searched the Global Biodiversity Information Facility (gbif.org) for occurrence points for all of the species in our study. We curated these results by removing duplicates, manually removing obvious outliers, and filtering out latitude and longitude coordinates that were precise to at least the third decimal place. This resulted in 657 occurrence points for *P. davidsonii*, 135 occurrence points for *P. rupicola*, 182 occurrence points for *P. fruticosus*, 115 occurrence points for *P. cardwellii*, and 1402 occurrence points for *P. newberryi*. We downloaded climate data from the WorldClim database (Hijmans, Cameron, Parra, Jones, & Jarvis, 2005) for the present-day, the mid-Holocene, and the LGM. Data for the current and mid-Holocene climates were at a resolution of 30 arc seconds, and LGM data were at a resolution of 2.5 minutes. Both the LGM and mid-Holocene data sets were generated by the CCSM4 model. We used only uncorrelated bioclimatic variables (Pearson’s *r* < 0.7), and used the same variables for each species. When given a decision about which correlated variable to remove, we retained variables that we suspected would be more important for explaining species’ distributions. The final variables used for SDM construction can be found in the Supplemental Table 2. After locality and climate data were curated, we built SDMs with the ensemble method implemented in the R package *biomod2* (Thuiller, Georges, Engler, Breiner, & Georges, 2016). We used four modeling approaches in *biomod2*: Random Forests, General Linear Models, Generalized Boosting Models, and Maximum Entropy as implemented in *Maxent* (Phillips, Anderson, & Schapire, 2006). We first cropped our raster files to −150° to −100° longitude and 35° to 65° latitude. We decided to use an extent that was larger than the range of our species of interest because it is plausible that species had distributions during the mid-Holocene and the LGM that are not confined within their current ranges. This extent also captures the distribution of the subgenus as a whole, including species that are not examined in this study. We ran five replicates per model, and randomly sampled 10,000 background pseudoabsences with the ‘random’ strategy in *biomod2*. We used 80% of our occurrence points for training models, 20% for testing models, and evaluated modules using the receiver operating characteristic (ROC) curve. For ensemble modeling, we only included models with an ROC score > 0.85, and weighted models based on their ROC score. Ensemble models were then forecast onto current, mid-Holocene, and LGM climate conditions.

### Models of demographic history and relationships between taxa

We used the R package *delimitR* (Smith & Carstens, 2020) to test models of demographic history within species. *delimitR* uses a machine learning algorithm to compare data simulated under demographic models of interest to a folded multi-dimensional site frequency spectrum (mSFS) constructed from high-throughput sequence data. Although *delimitR* was designed as software for species delimitation, it is able to compare models that differ with respect to the inclusion or absence of gene flow, divergence times, and relationships between lineages, making it a flexible and useful method to test models of demographic history more generally. Our main goal with *delimitR* was to determine whether intra-specific divergence occurred before or after the LGM. Specifically, our models focus on divergence times between lineages; did lineages diverge prior to the LGM, or did they diverge after the LGM? Model sets for both two-lineage and three-lineage species, as determined by STRUCTURE, can be seen in Figure 1. Our model set for species with two identified lineages includes the following scenarios: (1) post-LGM divergence (100 to 21,000 generations), (2) pre-LGM divergence (130,000 to 2,400,000 generations), and (3) pre-LGM divergence with secondary contact (migration). We defined pre-LGM divergence times to be between the start of the Pleistocene and the end of the interglacial cycle immediately preceding the LGM. Our reasoning for this is twofold. First, preliminary tests revealed that models were difficult to differentiate if priors on divergence times were too close to one another. Second, we suspect that the interglacial period immediately preceding the LGM would have had less suitable habitat (and thus species would have been the most isolated) than at any other point during the last glacial cycle (see Results). Any signal of divergence prior to the LGM should be captured with this broad prior space.

**Figure 1.**
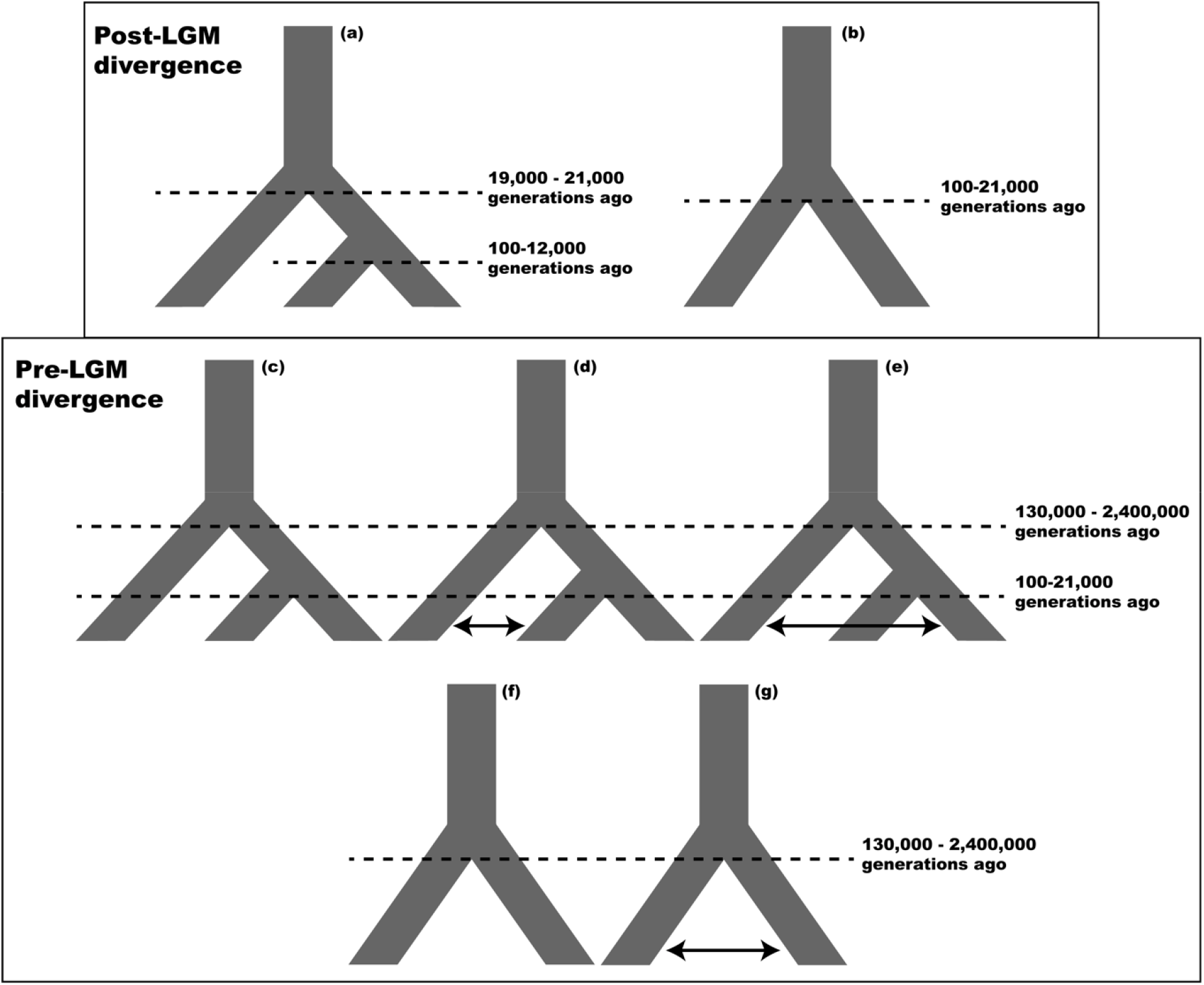
Demographic models implemented in *delimitR* for two-population (b, f-g) and three-population (a, c-e) scenarios. Two basic sets of models were tested: post-LGM divergence (a-b) and pre-LGM divergence (c-g). The two basic model sets differ in divergence time priors and the addition of secondary contact in pre-LGM divergence models.

Our model set for species with three lineages is an extension of the model set for two lineages. For post-LGM divergence, priors on divergence times correspond to the beginning of the Holocene to present day for the most recent divergence (100 to 12,000 generations ago), and to the estimated timing of the LGM ± 1,000 generations for the oldest divergence (19,000 to 21,000 generations ago). For pre-LGM divergences, priors on divergence times correspond to the LGM to the present day for the most recent divergence (100 to 21,000 generations ago), and to the beginning of the Pleistocene to the end of the last interglacial period for the oldest divergence (130,000 to 2,400,000 generations ago). We also included two pre-LGM divergence models with gene flow, allowing gene flow between each one of the sister lineages and the nonsister lineage (no gene flow between sister lineages). We tested this set of four models for all three possible topologies, for a combined total of twelve models. To conduct simulations in *delimitR*, we constructed five replicates of the mSFS with unlinked SNPs by randomly down-sampling 50% of the haplotypes assigned to each identified lineage with custom scripts available on the developer’s github (https://github.com/meganlsmith). We performed 20,000 simulations under each model in *fastsimcoa12* (Excoffier, Dupanloup, Huerta-Sánchez, Sousa, & Foll, 2013), and binned the mSFS for each species according to the 2N sample size of the population with the fewest samples, with a maximum of six bins. We used 500 decision trees to build the random forest classifier.

We used *SVDQuartets* (Chifman & Kubatko, 2014) to infer relationships between populations identified via STRUCTURE in a multispecies coalescent framework. For this analysis we used the data set which included all individuals across species boundaries, partitioned taxa by their identified genetic lineage, and set *Penstemon montanus* as the outgroup taxon. We evaluated all possible quartets and performed 100 bootstrap replicates.

### Hybridization and introgression between species

We used the software *HyDe* (Blischak, Chifman, Wolfe, & Kubatko, 2019) to detect instances of hybridization and introgression between focal taxa. *HyDe* uses phylogenetic invariants to detect hybridization between two parental lineages into a hybrid lineage, can differentiate between hybrid speciation and introgression, and can distinguish whether this gene exchange has occurred in a particular individual versus the population as a whole. We first ran a modified full *HyDe* analysis. To do so, we tested all possible combinations of parental and hybrid lineages, but because we are interested in gene flow between, rather than within species, we removed tests in which both parental lineages were from the same species. We also removed tests that included the Willamette population of *P. fruticosus*, as preliminary results indicated that this population produced spurious results, likely because of its small sample size (n = 10). This resulted in 423 total tests for interspecific hybridization. From there, we filtered out tests which did not produce statistically significant results at α = 0.05 after a Bonferroni correction. To assess the degree to which populations vs. individuals are introgressed, we subsequently implemented individual and bootstrap *HyDe* analyses on the parent-hybrid triplets that produced significant results from the modified full analysis.

## Results

### Data generation

For the combined data set, our filtering process resulted in 1,739 total retained loci, with an average of 1,502 loci per sample (±166), and 5,061 total parsimony-informative sites. Average heterozygosity was estimated to be 0.0105 ± 0.0077, and our data matrix had 13.73% missing SNPs. Results for both inter- and intra-specific data sets are summarized in Table 1.

**Table 1.** Post-processing GBS data generation statistics. All rows refer to intraspecific data sets, except for the final row, which refers to the interspecific data set. LPS = average loci per sample; TPI = total parsimony informative sites; *H_e_* = average heterozygosity estimates.

| Species | Total loci | LPS | TPI | $H_e$ | % missing SNPS |
| --- | --- | --- | --- | --- | --- |
| *P. rupicola* | 2454 | 2148.3 ± 235.01 | 2645 | 0.0106 ± 0.0094 | 13.11% |
| *P. cardwellii* | 2476 | 2192.3 ± 265.2 | 3649 | 0.0075 ± 0.0043 | 12.95% |
| *P. newberryi* | 2382 | 2129.1 ± 206.6 | 2159 | 0.0093 ± 0.0059 | 12.00% |
| *P. davidsonii* | 2494 | 2185.9 ± 243.6 | 3339 | 0.0144 ± 0.0085 | 12.89% |
| *P. fruticosus* | 3333 | 2656.6 ± 388.3 | 2128 | 0.0104 ± 0.0092 | 19.08% |
| Total (inter-species) | 1739 | 1502.5 ± 165.9 | 5061 | 0.0105 ± 0.0077 | 13.74% |

### Identification of populations

For *P. fruticosus, P. davidsonii*, and *P. newberryi*, we used K = 2 for downstream analyses, and for *P. rupicola* and *P. cardwellii*, we used K = 3 (Figure 2). Explanations of our decisions for each species can be found accompanying Supplemental Figure 1. K values for *P. fruticosus* correspond loosely to one cluster representing the northern extent of the species’ range in the Cascades and into Ochoco National Forest, and the other cluster representing the southern extent of the species’ range in the Cascades. For *P. rupicola*, a north-south gradient is evident. The first of the three identified cluster for this species corresponds to the southern-most extent of the species’ range in the Klamath Mountains. The second cluster corresponds to the southernmost extent of the Cascades, Mt. Shasta, the Klamath Mountains in southeastern Oregon, and near Crater Lake. The third cluster corresponds to the northern-most extent of the species’ range in the north Cascades, and extends south to Sisters Mountains in Oregon. There is a clear grade between the northern and southern clusters that begins near Crater Lake and ends near the Columbia River. There is also an outlier introgressed location north of the Columbia River, which appears to be admixed between all three clusters. For *P. davidsonii*, one cluster is found almost exclusively north of the Columbia River, and the other cluster is exclusively south of the Columbia River, with admixture in two locations near Crater Lake and near Mt. Jefferson. For *P. cardwellii*, one cluster corresponds to the southern-most extent of the species’ range, in the Klamath Mountains. A second cluster corresponds to the coast range north of the Klamath Mountains, and into the Cascades near Mt. St. Helens and just south of the Columbia River. The third cluster corresponds to the eastern portion of this species’ range, beginning near Mt. Hood and moving southwards along the Cascades crest.

**Figure 2.**
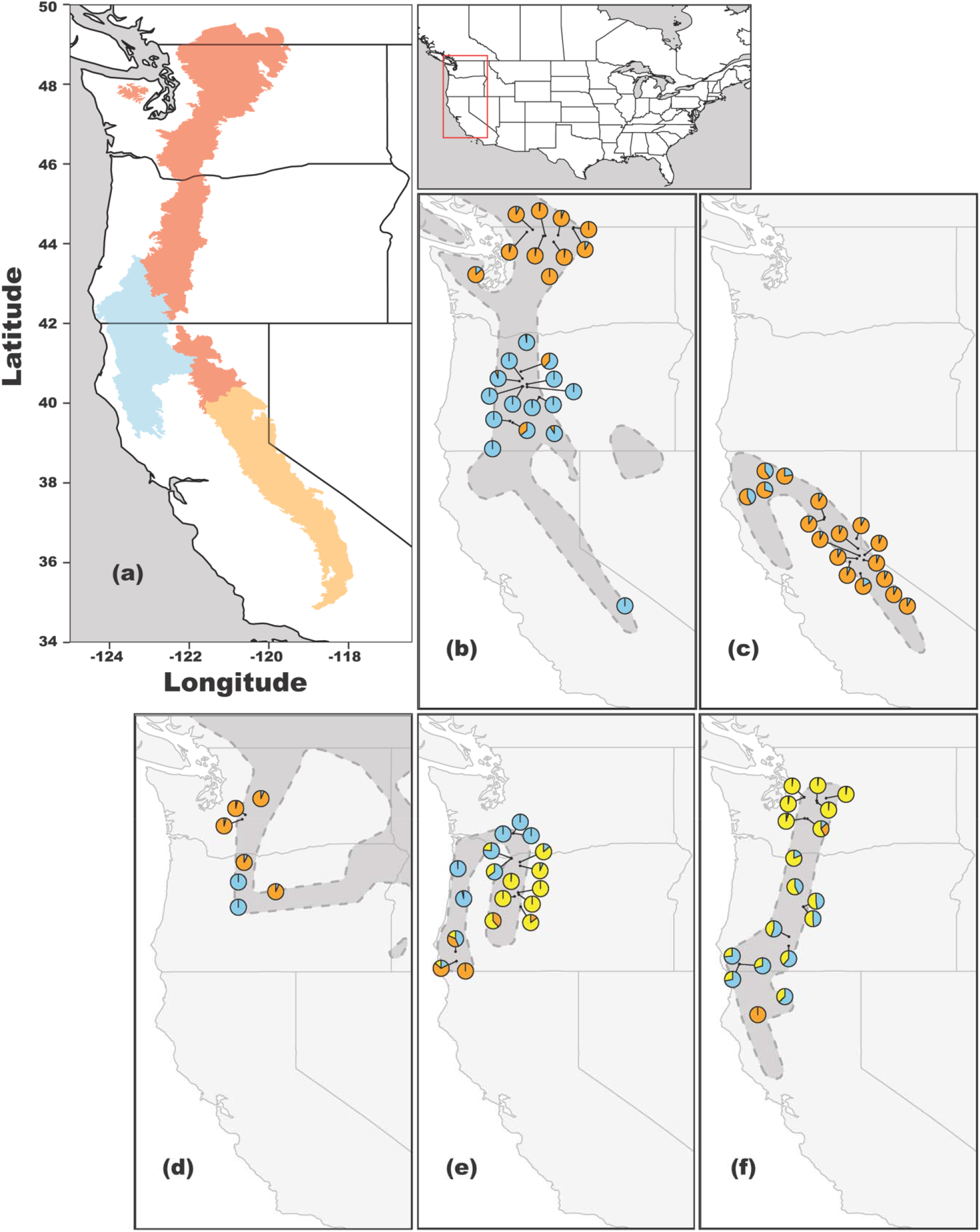
(a) Map of the Pacific Northwest. Highlighted areas correspond to the Sierra Nevada (orange), Cascades and North Cascades (red), and the Klamath Mountains (blue), as defined by the United States Environmental Protection Agency. (b-f) STRUCTURE results plotted on collection localities for (b) *P. davidsonii*, (c) *P. newberryi*, (c) *P. fruticosus*, (e) *P. cardwelii*, and (f) *P. rupicola*. Colors in the pie charts correspond to the probability of membership of an individual to each of the *K* intraspecific populations. Colors for one species do not correspond to colors in a different species, and darkly shaded areas represent approximate species’ ranges.

### Species distribution models

Average ROC scores across replicates and modelling strategies were generally high. General results for each species are reported in Table 2, and average variable importance across replicates for each model and variable are reported in Supplemental Table 2. SDMs for the present day identified suitable habitat that extends beyond the known natural range of three species (*P. rupicola, P. fruticosus*, and *P. newberryi*) (Figure 3). In addition, for all species, suitable habitat shifted north during the mid-Holocene warm period (Figure 3). Conversely, there was a distinct southward shift in suitable habitat for three species (*P. cardwellii, P. newberryi, P. rupicola*) during the LGM, and there was noticeably more suitable habitat for all species but one (*P. cardwellii*) during this time (Figure 3).

**Figure 3.**
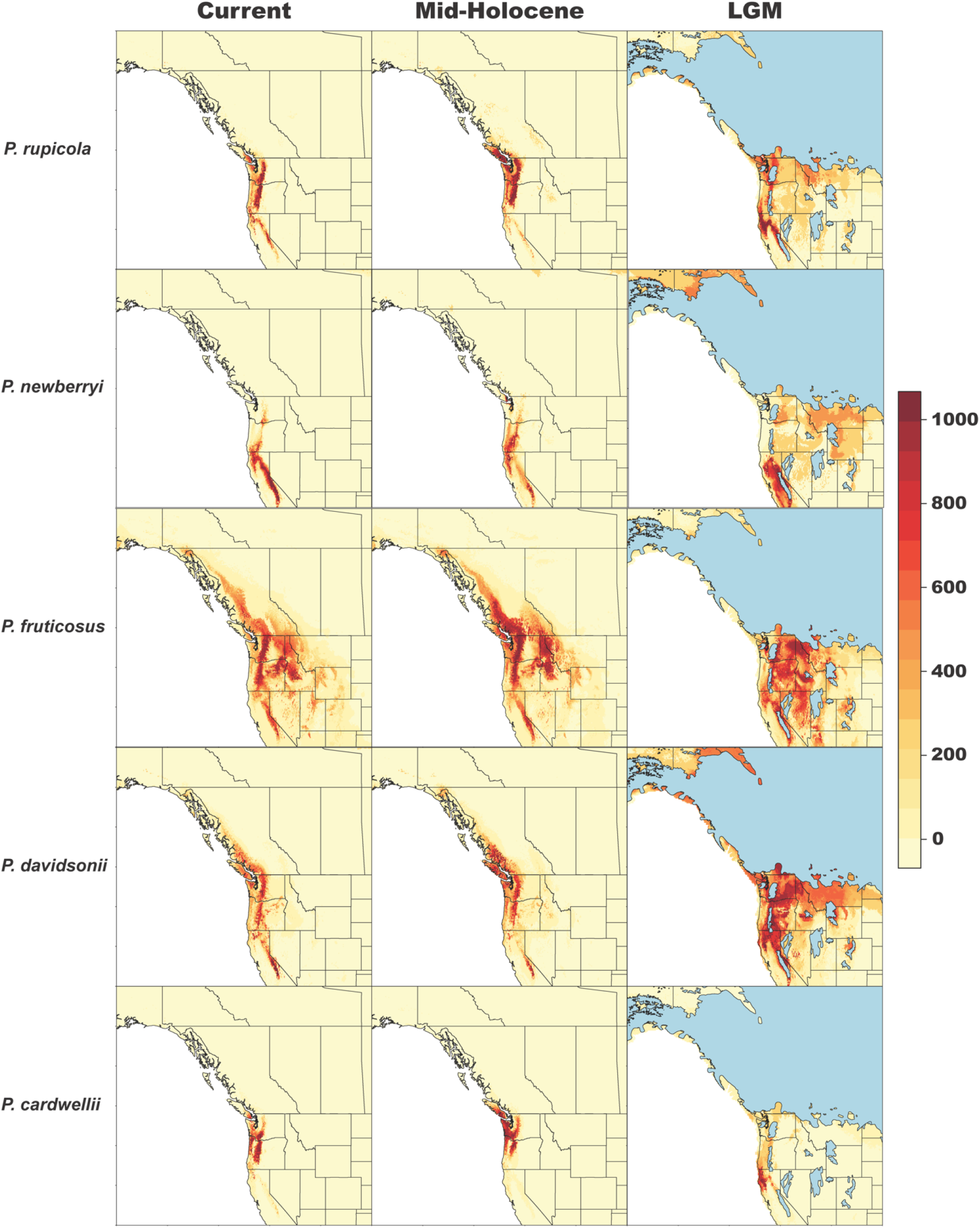
Species distribution models (SDMs) for five *Penstemon* subgenus *Dasanthera* species. From left to right, plots indicate projections for the present-day, the mid-Holocene warm period, and the LGM. Habitat suitability scores are represented by the heat map on the right; warmer colors indicate higher habitat suitability. The light blue regions in the LGM plots represent areas with glacial cover.

**Table 2.** Results of species distribution modeling. Numbers correspond to the average of ROC values across 5 replicates for each model. Models correspond to the Maximum Entropy model as implemented in *Maxent* (MAXENT.Phillips), General Linear Models (GLM), Random Forests (RF), and Generalized Boosting Models (GBM).

| Species | MAXENT.Phillips | GLM | RF | GBM |
| --- | --- | --- | --- | --- |
| *P. rupicola* | 0.941 | 0.994 | 0.996 | 0.995 |
| *P. cardwellii* | 0.986 | 0.991 | 0.997 | 0.983 |
| *P. newberryi* | 0.965 | 0.997 | 0.997 | 0.997 |
| *P. davidsonii* | 0.874 | 0.992 | 0.994 | 0.994 |
| *P. fruticosus* | 0.957 | 0.955 | 0.968 | 0.960 |

### Models of demographic history

We found support for post-LGM divergence models for three out of five species (Figure 4; Table 3). We also found support for a pre-LGM divergence model for one species (*P. rupicola*), and were unable to reach any definitive conclusion for another (*P. fruticosus*). For the three species supported by post-LGM models (*P. cardwellii, P. newberryi, P. davidsonii*), average error rates were low, at a rate near or below 0.05 (Table 3). However, for the two remaining model sets, average error rates were moderate-to-high, at 0.192 (*P. rupicola*) and 0.202 (*P. fruticosus*) Average posterior probabilities tended to be the highest for species with low error rates, and *vice versa*. For the two species with three-lineage models, (*P. cardwellii* and *P. rupicola*), lineages in the Klamath Mountains were more differentiated than the other two identified intraspecific lineages.

**Figure 4.**
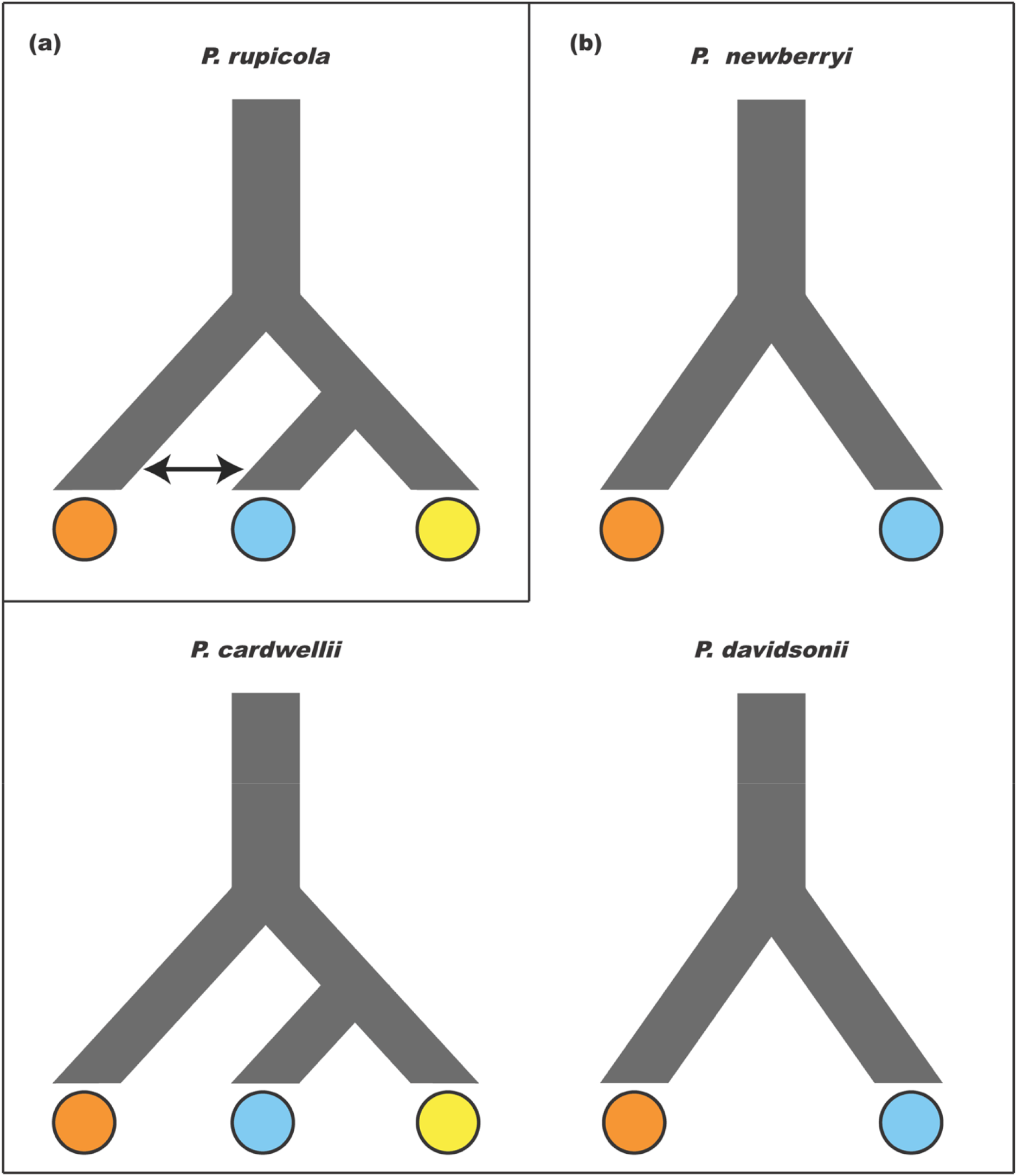
The most strongly supported demographic model for each species. No model for *P. fruticosus* is shown because of the equivocal results for that species. Colors correspond to the clusters shown in Figure 2. Models are separated on the basis of (a) pre-LGM and (b) post-LGM divergence.

**Table 3.** Results of the demographic model tests in *delimitR* for each species. Average error and average posterior probabilities correspond to the average of the out-of-bag error rates and posterior probabilities across five replicates.

| Species | Average Error Rate | Best Model | Average Posterior Probability |
| --- | --- | --- | --- |
| *P. rupicola* | 0.192 | Pre-LGM divergence + migration | 0.611 |
| *P. cardwellii* | 0.043 | Post-LGM divergence | 0.987 |
| *P. newberryi* | 0.048 | Post-LGM divergence | 0.939 |
| *P. davidsonii* | 0.059 | Post-LGM divergence | 0.995 |
| *P. fruticosus* | 0.202 | Equivocal | 0.452 |

*Penstemon rupicola* is the only species that strongly supported a pre-LGM divergence model. In all replicates, the best model for this species was one of pre-LGM divergence, had the northern and southern lineages as sister, and included gene flow between the Klamath lineage and one of the other lineages (Figure 4). However, there was some uncertainty in model selection for this species. Posterior probabilities for the best model were low (Table 3), and average error rates for the best model were higher than any other model (Supplemental Table 3). The confusion matrix indicates that models identical in all aspects but their gene flow parameters (*e.g*., models 4, 5, and 6) were the most difficult to differentiate (Supplemental Table 3). Only one other model (model 1) received votes; however, this model is one of post-LGM divergence, and does not include gene flow, although it has the same topology as models 4-6. We interpret these results as strong support for the relationships between *P. rupicola* lineages, and a reasonable degree of support for the oldest divergence times between populations preceding the LGM.

A post-LGM divergence model was supported for three species: *P. cardwellii, P. newberryi*, and *P. davidsonii*. For *P. cardwellii*, the best model was one of post-LGM divergence, with the eastern and western populations sister to one another (Figure 4). No pre-LGM divergence model received more than 5% of the votes (Supplemental Table 4). For *P. newberryi*, all replicates strongly supported the post-LGM divergence model (Figure 4; Table 3). The only other model that received votes for *P. newberryi* was the pre-LGM divergence that included gene flow (Supplemental Table 5). The same is true for *P. davidsonii;* all replicate runs for *P. davidsonii* strongly supported the post-LGM divergence model, and the vast majority of votes given to alternative models were given to the pre-LGM divergence model with secondary contact (Supplemental Table 6).

Demographic model tests for *P. fruticosus* were inconclusive, as error rates were high, and posterior probabilities were low (Table 3). Each of the three models received the most votes in at least one replicate (Supplemental Table 7). We therefore cannot determine the best demographic scenario for *P. fruticosus*.

### Relationships between lineages

The lineage tree inferred with *SVDQuartets* presents an overall strongly supported topology (Figure 5). Immediately apparent is the strong geographic pattern present, especially with respect to the Klamath Mountains, as all of the Klamath lineages from separate species form a single clade with strong support (100% bootstrap). Second, *P. cardwellii, P. fruticosus*, and *P. newberryi*, as currently circumscribed, are polyphyletic. Of the three identified genetic lineages of *P. cardwellii*, none are sister to one another. *Penstemon newberryi* var. *berryi* is sister to the Klamath lineage of *P. rupicola*, although this pair is then sister to *P. newberryi* var. *newberryi*. The northern lineage of *P. fruticosus* is sister to *P. davidsonii* with high bootstrap support (100%), which the southern lineage falls within the clade containing the other four species, and is sister to the eastern lineage of *P. cardwellii*.

**Figure 5.**
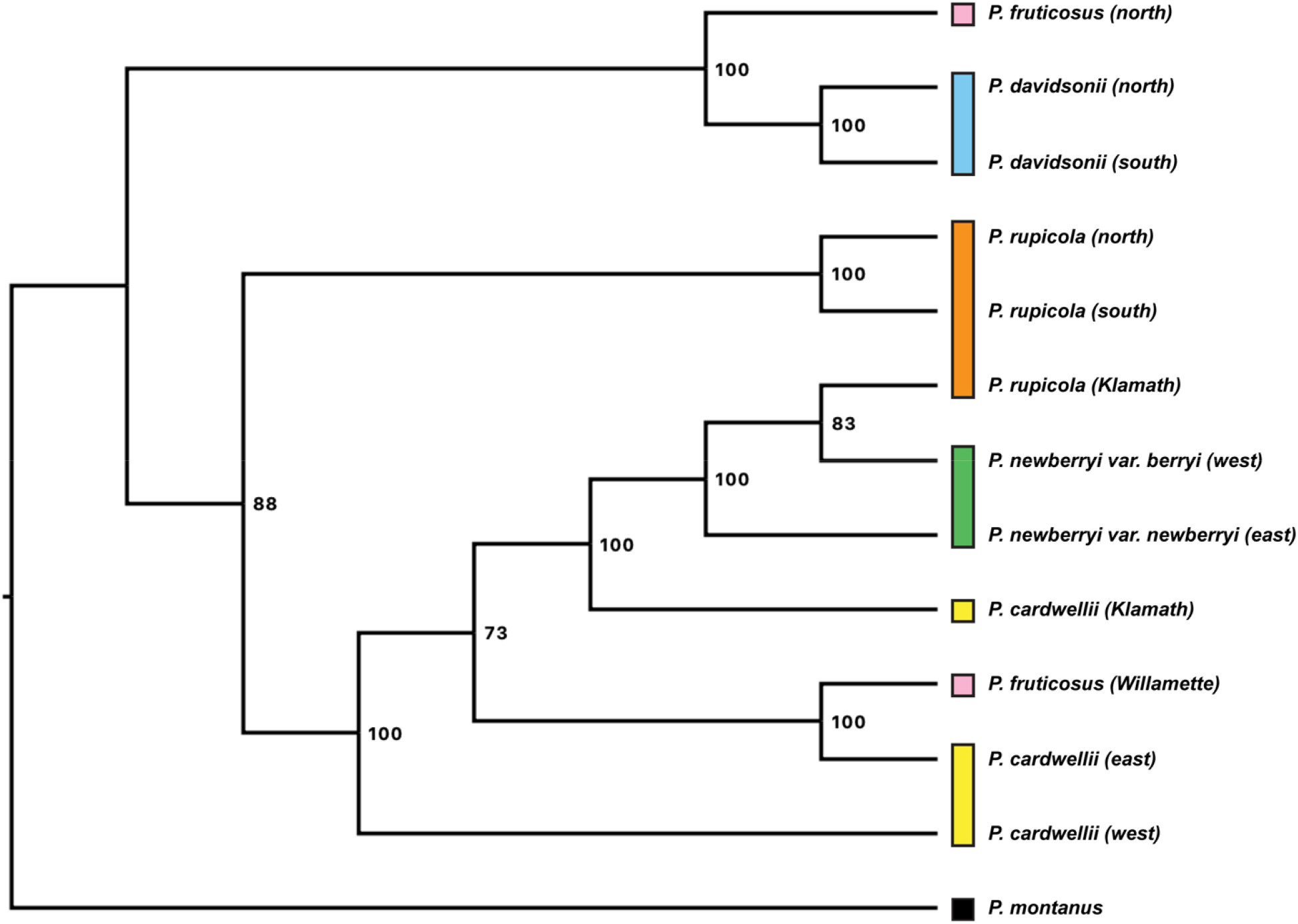
Lineage tree constructed with *SVDQuartets*. Bootstrap scores are presented at the nodes. An individual’s lineage was determined by the majority genetic cluster for that individual as inferred by the *STRUCTURE* analyses. Each color corresponds to a nominal species.

### Hybridization and introgression between species

Our tests of hybridization and introgression revealed substantial evidence for hybridization between species. Of the 423 total tests considered, 45 of these (10.6%) were statistically significant (Supplemental Table 8). Of these significant results, 39 of them (86.7%) included at least one lineage from the Klamath Mountains, either as a parent or a hybrid lineage. Since only 273 of the initial 423 tests (64.5%) matched this criterion, lineages from the Klamath Mountains are overrepresented as probable players in hybridization in *Dasanthera*. This overrepresentation of lineages from the Klamath Mountains is visualized in a heat map in Supplemental Figure 2. Here, the Klamath lineages are clearly involved in more putative hybridization events than any other lineages, both between themselves and between non-Klamath lineages. Further inspection of putative hybridization events involving no Klamath lineages identified (1) the ‘southern’ *P. davidsonii* lineage and (2) the ‘southern’ *P. rupicola* lineage as potential hybrid lineages.

There are four putative parental sources responsible for the formation of the southern *P. davidsonii* lineage: the northern *P. davidsonii* lineage, the eastern *P. cardwellii* lineage, the southern *P. rupicola* lineage, and the northern *P. fruticosus* lineage. The distribution of *γ* across bootstrap replicates for each putative hybridization event can be found in Supplemental Figure 3a. Our results indicate that the putative hybridization events forming the southern *P. davidsonii* lineage likely represent instances of introgression, rather than hybrid speciation, since average *γ* values are not close to 0.5. Further, since we do not see a pattern of jumping between *γ* values with no estimates between, these results likely indicate uniform admixture (Blischak *et al*., 2018).

Like the southern *P. davidsonii* lineage, our analyses with *HyDe* identified four parental sources putatively responsible for the formation of the southern *P. rupicola* lineage. All combinations include the northern *P. rupicola* lineage and either the eastern or northern *P. cardwellii* lineages, or the southern *P. davidsonii* lineage. The distribution of *γ* across bootstrap replicates (Supplemental Figure 3b) suggests that, again, similar to the southern *P. davidsonii* lineage, these hybridization events likely represent introgression rather than hybrid speciation, and admixture is likely uniform.

Generally, our results for non-Klamath lineages suggest introgression, rather than hybrid speciation, as the mechanism by which *Dasanthera* species exchange genes. The reasons for this are potentially twofold: (1) we intentionally did not include obvious hybrids, focusing our efforts on the demographic history of species and understanding hybridization in a phylogenetic context, and (2) our sampling is broad, rather than at the population scale, meaning we are considering lineages at a phylogenetic scale rather than at a population-genetic scale.

## Discussion

We examined population structure, inferred relationships between species, modeled demographic histories, and reconstructed species distribution models for five PNW-distributed species in *Penstemon* subgenus *Dasanthera*. Our analyses suggest that Quaternary glacial cycles played a key role in shaping species’ distributions through time, which in turn affected patterns of intraspecific genetic diversity. We uncovered a prevalent north-to-south axis of genetic differentiation that is consistent with patterns observed in other taxa from the region (*e.g*., Soltis *et al*., 1997). In addition, we find evidence that the bulk of intraspecific genetic variation in these species is likely not due to vicariance prior to the LGM (Figure 4). To the contrary, our SDMs suggest that most species’ ranges expanded and experienced greater connectivity during the LGM than after glacial retreat (Figure 3). Of particular phylogeographic importance for these species are the Klamath Mountains, as they host a large portion of genetic diversity (Figure 2), and appear to be a hotspot for gene exchange both within and between species (Supplemental Figure 2). In turn, the large degree of hybridization in the Klamath Mountains blurs species limits and makes inferring relationships between species in this region challenging (Figure 5).

### The near-absence of glacial refugia

We posit that for *P. newberryi, P. fruticosus*, and *P. davidsonii*, large amounts of suitable habitat during the LGM resulted in greater population connectivity, facilitating both intra- and inter-specific gene flow. Subsequent range reductions during the mid-Holocene then caused population fragmentation, resulting in species’ distributions that are similar to the present day. In turn, this geographic isolation may have led to population-genetic substructure, which was then identified with our *STRUCTURE* analyses. *Penstemon cardwellii* is the most mesic-tolerant *Dasanthera* member, is found in both the Oregon Coast Range as well as the inland Cascades, and was the only species in this study with less suitable habitat during the LGM. Consequently, coastal *P. cardwellii* populations may have exhibited a response to glaciation (Figure 3) similar to other coastal species associated with PNW mesic flora (*e.g*., Smith *et al*., 2018). Coupled with the distinct reduction in suitable habitat for *P. cardwellii* is a southern range shift along the Pacific coast. There is little suitable habitat north of the Klamath Mountains for this species, implicating this region as a potential LGM refugium. *Penstemon rupicola*, like most of the other species, likely shifted its range south during the LGM (Figure 3). However, this species is the only one that supported a model of pre-LGM divergence (Figure 4), suggesting that the *P. rupicola* lineage from the Klamath Mountains may have been isolated since the last interglacial period, or even earlier. An important caveat to consider is that while many of these species appear to exhibit a signal of recent (post-LGM) divergence, this signal does not preclude - and in fact likely obscures - more ancient and complex demographic histories. The PNW has experienced repeated glaciation events throughout the Pleistocene, all of which undoubtedly altered species’ distributions and affected the geographic context in which gene flow occurred (Hewitt, 2004; Shafer *et al*., 2010). Given the age of the *Dasanthera* clade, estimated to have formed during the early Pleistocene roughly 1.9 MYA (Wolfe *et al*., unpublished data), it is likely that all of the species examined in this study would have experienced several oscillations of colder periods during glaciation and warmer periods following glacial retreat. The simple models of demographic history that we have employed in this study are, by virtue of their design, unable to capture that complexity.

The classic paradigm regarding species’ responses to glacial cycles posits that, generally, temperate species will contract their ranges during peak glacial activity, often congregating in refugia, and subsequently expand their ranges during interglacial periods (Hewitt, 2000; Hewitt, 2004). However, for four of the five taxa examined here, we observe the inverse pattern; it appears that most *Dasanthera* species expanded their ranges during the LGM, and subsequently retracted their ranges in the ensuing interglacial period (Figure 3). While this pattern has been observed in other species, typically it has been restricted to cold-adapted taxa (Martinet *et al*., 2018; Stewart *et al*., 2010) or to Neotropical systems (Leite *et al*., 2016; Perez, Bonatelli, Moraes, & Carstens, 2016), although see Gür (2013). Our results suggest that the longstanding consensus of temperate species’ responses to glacial cycles may not be as generally applicable as previously thought.

### The ‘central’ importance of the Klamath Mountains

Our analyses suggest the presence of suitable habitat in the Klamath Mountains during the LGM for every species included in this study (Figure 3). This, combined with our analyses implicating this region as a hotspot for interspecific hybridization (Supplemental Figure 2) and genetic differentiation (Figure 2), highlight the phylogeographic importance of the Klamath Mountains to *Dasanthera* species. As noted earlier, this region has been identified as an important geographic feature for many plant and animal taxa (Eckert *et al*., 2008; Furnier & Adams, 1986; Goebel *et al*., 2009; Gugger *et al*., 2010; Kiefer *et al*., 2009; Kuchta & Tan, 2005; Smith &Sawyer, 1988; Soltis *et al*., 1997; Patterson & Givnish, 2003; Pelletier *et al*., 2011; Whittaker, 1961). In particular, the Klamath region is thought to owe much of its diversity to its topographical complexity and its proximity to several other mountain ranges, including the Cascades and Sierra Nevada Mountains, and the coastal ranges of Oregon and California. This allows species from more northern latitudes to access the area when the climate cools, and then persist at higher elevations when conditions warm again (Smith & Sawyer, 1988; Whittaker, 1961). We hypothesize that the Klamath Mountains have served a similar role for *Dasanthera* species, and suggest that this region serves as a ‘choke-point’ for species’ movement between the Cascades and Sierra Nevada Mountains. The following scenario could explain both the abundance of *Dasanthera* diversity in the Klamath Mountains and the relative lack of diversity in the Sierra Nevada Mountains. (1) As the climate cools, species distributed at more northerly latitudes shift their ranges southward. (2) Species’ distributions begin to experience more overlap, and the poor long-distance dispersal ability of *Dasanthera* leads to species’ ranges overlapping in the Klamath Mountains, as it is the only geographically proximate area with suitable habitat. (3) Species begin to exchange genes when in sympatry, forming complex hybridization networks and blurring species limits in this region. The resulting hybrids that persist form hybrid swarms (lineage fusion), which, over time, become the predominant forms in the region. (4) These hybrids become less fit and more susceptible to genetic homogenization from parental taxa the further from the Klamath Mountains they get, until they grade into *P. davidsonii/P. newberryi* in the Sierra Nevada Mountains, or *P. davidsonii/P. rupicola* in the Cascades.

### Interplay between hybridization and species limits in Penstemon subgenus Dasanthera

Hybridization has undoubtedly made identifying relationships between *Dasanthera* species challenging. Previous efforts have been conducted to infer relationships between species using nuclear and chloroplast sequence data, and inter-simple sequence repeat markers (Datwyler & Wolfe, 2004). The tree produced in Datwyler and Wolfe (2004) suffers from low bootstrap support along the backbone of the tree, and its topology differs substantially from the tree presented in this study (Figure 5). This is due, at least in part, to the relative lack of parsimony-informative sites in the *ITS* and *matK* sequence data, but the prevalence of hybridization across this subgenus also almost certainly contributed to the uncertainty in species relationships observed in Datwyler and Wolfe (2004). It is worth noting that Albert Every, using morphological and chemical features of *Dasanthera* taxa, suggested the same relationships between species in subgenus *Dasanthera* as identified in this study (Every, 1977). Every (1977) also noted the abundance of gene flow between species in the Klamath region, and suggested a complicated network where *P. newberryi* var. *berryi* was freely exchanging genes with *P. rupicola* and *P. cardwellii*, and that genes from *P. cardwellii* were introgressed into *P. rupicola*, which in turn introgressed into *P. davidsonii*. While the exact details of Every’s hypotheses were not explicitly tested here, they do serve as a useful indicator of the complex demographic history of *Dasanthera* in the Klamath Mountains.

Despite our findings, species limits in *Dasanthera* will need revisited, at least in the context of the clade containing *P. cardwellii* and the lineages from the Klamath Mountains. The work presented here elucidates some of the relationships among species distributed in the Cascades and Sierra Nevada Mountains with strong support. However, the prevalence of gene flow between species may confound these inferences and provide a false signal of confidence. Because we have shown that the evolutionary history of these species includes introgression events, species’ relationships are likely more accurately portrayed as a network; any evolutionary history of *Dasanthera* species depicting solely bifurcating lineages is therefore an insufficient explanation of the relationships between species in this subgenus. As a result, the tree presented in this work should be interpreted with caution, especially for lineages in the Klamath Mountains.

While we have explored the occurrence and geographic location of hybridization, the mechanisms underlying why it occurs – or at least, the reason why hybrids appear to persist – remains unanswered. Conversely, the question remains: what maintains species boundaries between *Dasanthera* species at all? Cross-fertilization experiments have verified that there are likely few cytogenetic barriers, if any, that prevent the formation of hybrid taxa (Every, 1977; Viehmeyer, 1958). Several studies focusing on hybrids between *P. newberryi* and *P. davidsonii* have supported this finding, and have elaborated on questions regarding hybrid fitness (Clausen *et al*., 1940; Kimball, 2008; Kimball, Campbell, & Lessin, 2008; Kimball & Campbell, 2009). *Penstemon newberryi* and *P. davidsonii* are the only two *Dasanthera* species located in the Sierra Nevada Mountains, and they form extensive hybrid zones where their ranges overlap (Clausen *et al*., 1940; Every, 1977). Investigations into the mechanisms controlling hybrid formation and persistence uncovered that hybrids are likely formed due to a shared pollinator community (Kimball, 2008), and that intermediate resource use and physiological tolerances likely allow hybrids to persist in intermediate environments (Kimball & Campbell, 2009). There also appear to be few cytogenetic barriers to hybrid formation between these species, although there is an apparent maternal effect to hybrid fitness with respect to elevation (Kimball *et al*., 2008).

Despite this, we have uncovered extensive hybridization between other *Dasanthera* species in the Klamath Mountains, the genetic signatures of which extend well beyond this geographic region into source populations of parental species. Furthermore, we identified two additional lineages (southern *P. davidsonii* and southern *P. rupicola*) not associated with Klamath Mountain lineages that exhibit evidence of introgression from other taxa (Supplemental Figure 3). The widespread persistence of this introgression implies that F1 hybrids are able to backcross into their parental species, making genetic differentiation via reinforcement unlikely. Therefore, while there is evidence suggesting that reduced hybrid fitness can maintain species boundaries via reinforcement in *P. davidsonii* x *P. newberryi* hybrid zones, our results indicate that reinforcement is likely not maintaining species boundaries in most other *Dasanthera* hybrid zones. In these cases, it seems more probable that species boundaries are formed and maintained when species’ ranges do not overlap, limiting gene flow due to poor dispersal ability.

## Acknowledgements

Funding was provided by the US National Science Foundation grant DEB-1455399. We thank the University of Washington Herbarium at the Burke Museum and the Oregon State University Herbarium for providing sample loans. We thank the Unity computing cluster at Ohio State University for the use of computing nodes for data processing and analysis. We thank Connor Lang for assistance with collections in the field, Dr. Shannon Datwyler for the donation of samples, and Dr. Megan Smith for advice on the GBS protocol. We also thank Dr. Megan Smith and members of the Wolfe lab for comments that improved this manuscript prior to publication.

## Data Accessibility

All scripts and parameter files used for data processing and analysis, along with unprocessed sequence reads and processed inter- and intraspecific reads, are available on Dryad (https://doi.org/10.5061/dryad.n5tb2rbtf).

## Author Contributions

B.W.S and A.D.W designed the study and collected samples. B.W.S collected genomic data, performed analyses and wrote the manuscript. B.W.S and A.D.W edited the manuscript and approved its final version.

