## supplement for "Phylogeographic analysis of shrubby beardtongues reveals range expansions during the Last Glacial Maximum and implicates the Klamath Mountains as a hotspot for hybridization"

**Table of Contents:**

| **Supplemental Figure 1** | Page 2 |
| --- | --- |
| **Supplemental Figure 2** | Page 3 |
| **Supplemental Figure 3** | Page 4 |
| **Supplemental Table 1** | Pages 5-6 |
| **Supplemental Table 2** | Page 7 |
| **Supplemental Table 3** | Page 8 |
| **Supplemental Table 4** | Page 9 |
| **Supplemental Table 5** | Page 10 |
| **Supplemental Table 6** | Page 11 |
| **Supplemental Table 7** | Page 12 |
| **Supplemental Table 8** | Page 13 |

*Supplemental Figure 1*. ∆K *STRUCTURE* plots (bottom, left) and likelihood plots (bottom, right) for each species. Our justifications for choosing K values for each species are as follows:

*Penstemon rupicola*. ∆K suggests K = 2, followed by K = 3. However, there is a jump in likelihood from K = 2 to K = 3, and K = 3 is near the apex of likelihood estimates for *P. rupicola*. We therefore chose K = 3 for *P. rupicola*.

*Penstemon fruticosus.* ∆K suggests K = 2, and the likelihood for this value is not much different than for K = 3 (the next best K value). Observation of the K = 3 plots only incorporates a ‘ghost’ third population in roughly equal proportions for all individuals. For this reason, and given the low sample size for this species, we used K = 2 for *P. fruticosus*.

*Penstemon davidsonii.* ∆K suggests K = 6, but observations of the likelihood plot indicate that this is an artifact caused by the dramatic decline in likelihood for K = 7. We therefore chose the second best ∆K value for *P. davidsonii*, K = 2.

*Penstemon newberryi.* ∆K suggests K = 3, but observations of the STRUCTURE plots for this species for any K value larger than K = 2 only adds ‘ghost’ populations, in roughly equal proportions, to all individuals. At K = 2, the distinction between populations corresponds to the two varieties of *P. newberryi*, with populations that possess both genetic lineages occurring almost exclusively in the Klamath Mountains. It seems likely that the genetic differentiation in this species reflects a grade into a hybrid taxon (*P. newberryi* var. *berryi*). So, while all individuals at K = 2 belong by majority to the same population, we separated individuals based on their subspecific distinction. We did so to explore the genetic diversity that is present in this species, even if this separation does not correspond to wholly different populations *per se*.

*Penstemon cardwellii.* ∆K suggests K = 2 for this species, but the likelihood plot increases until K = 5. Geographic subdivision was evident between populations up to K = 5. We decided to use K = 3 for *P. cardwellii* because it describes most of the genetic differentiation between populations while also maintaining a reasonable number of models to test with *delimitR*.

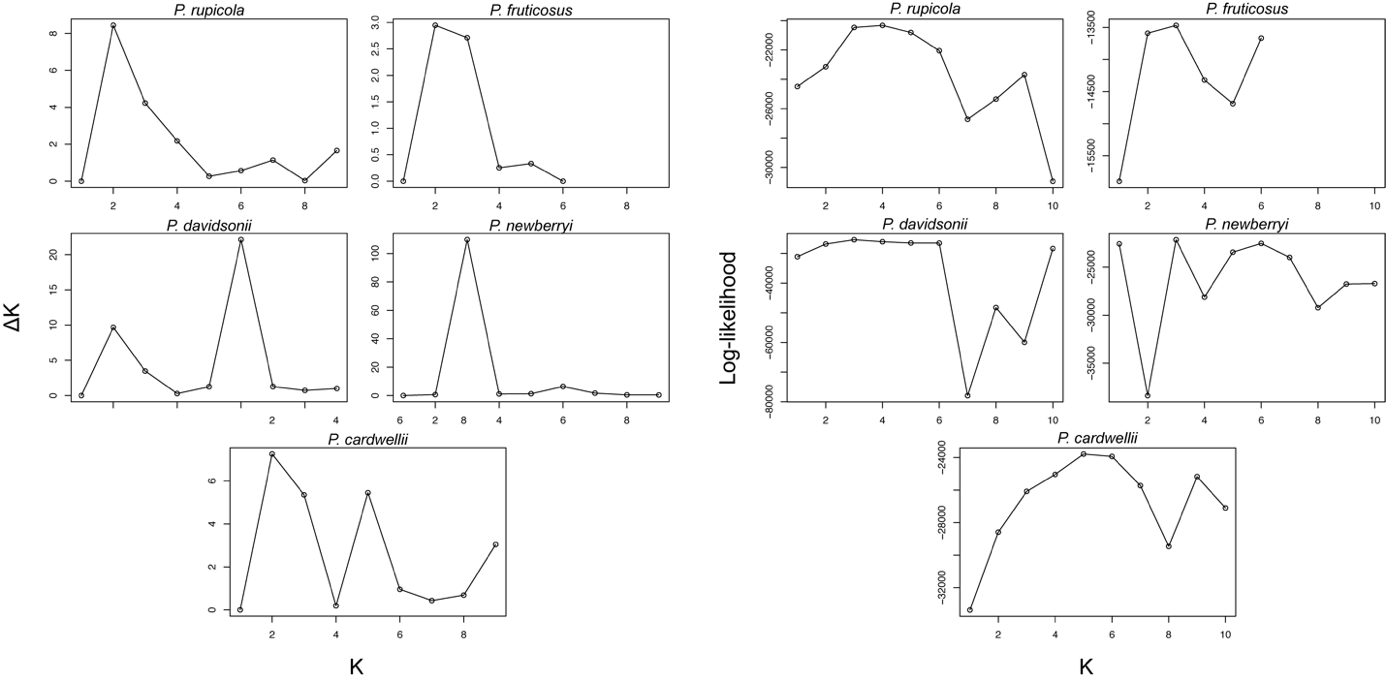

*Supplemental Figure 2.* Heatmap of significant *HyDe* results. Warmer values indicate more interactions, either as a parent or a hybrid. Values are calculated as the proportion of tests involving two lineages that are statistically significant. Lineages from the Klamath Mountains are outlined in black.

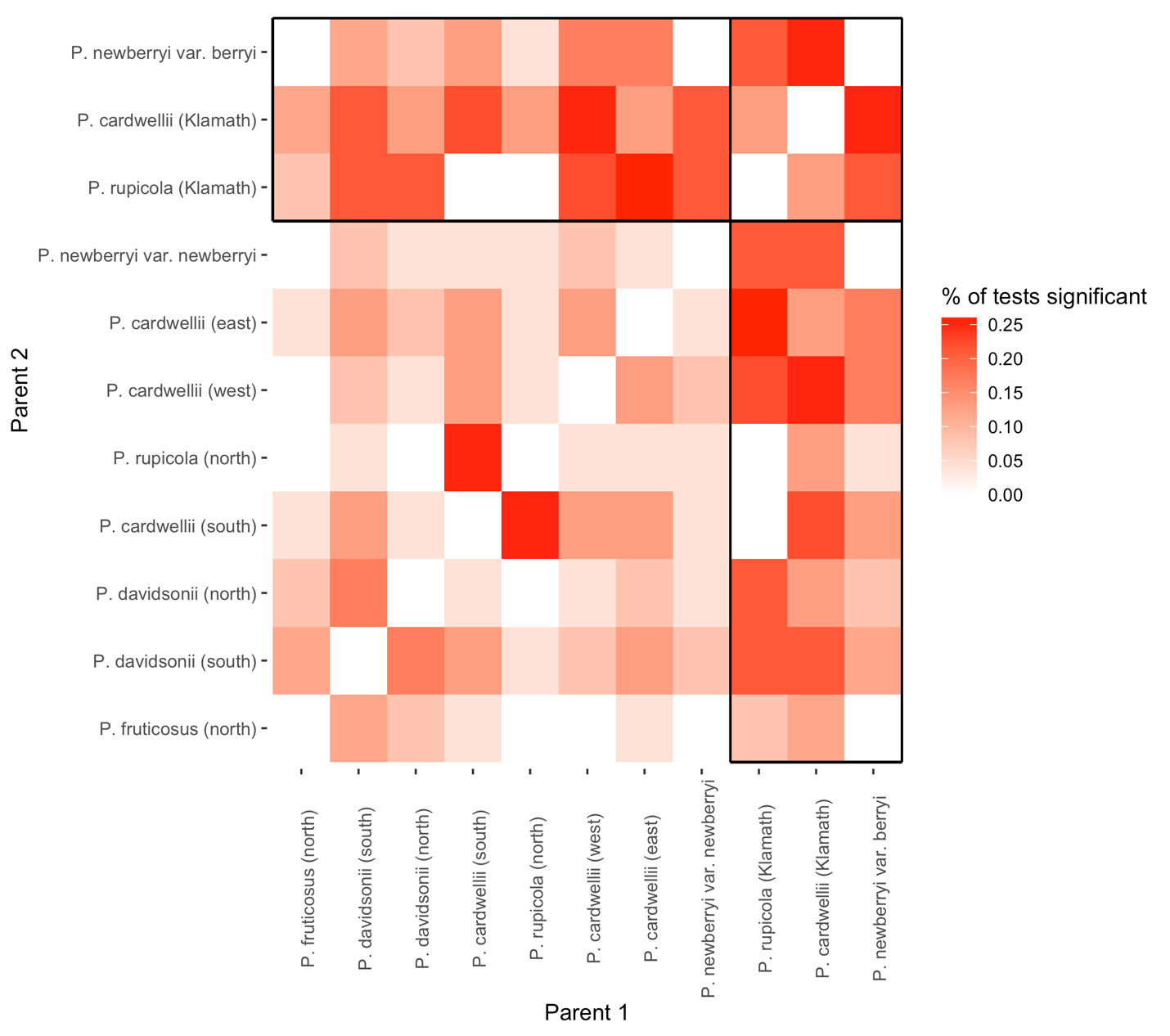

*Supplemental Figure 3.* Violin plots of 𝛾 across 100 bootstrap replicates for the southern *P. davidsonii* lineage (a) and the southern *P. rupicola* lineage (b).

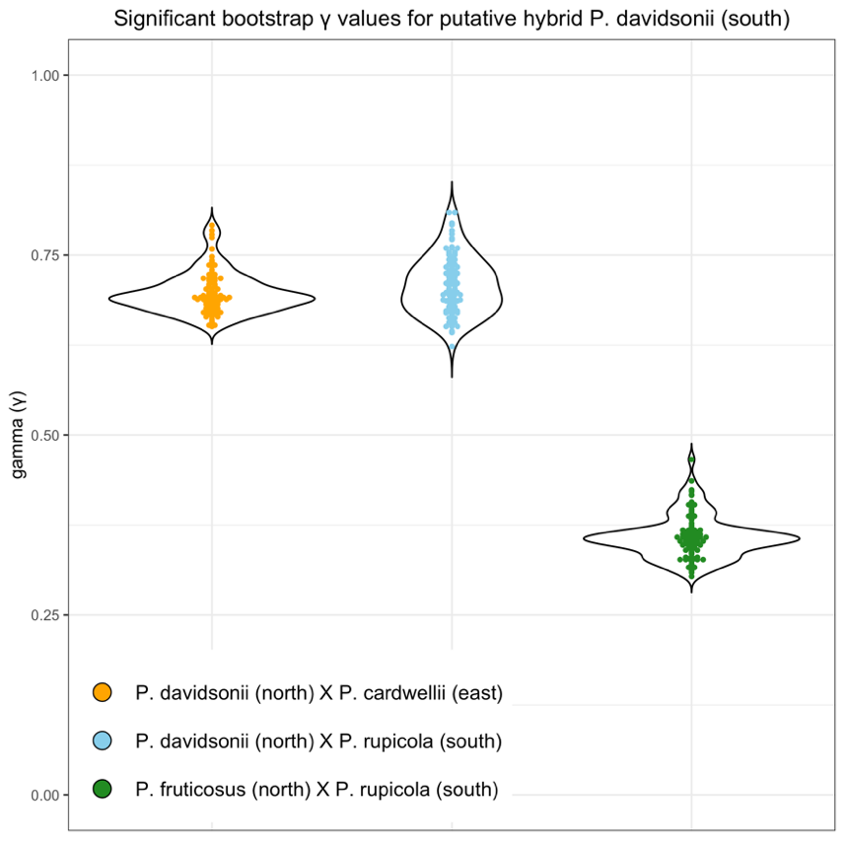

(a)

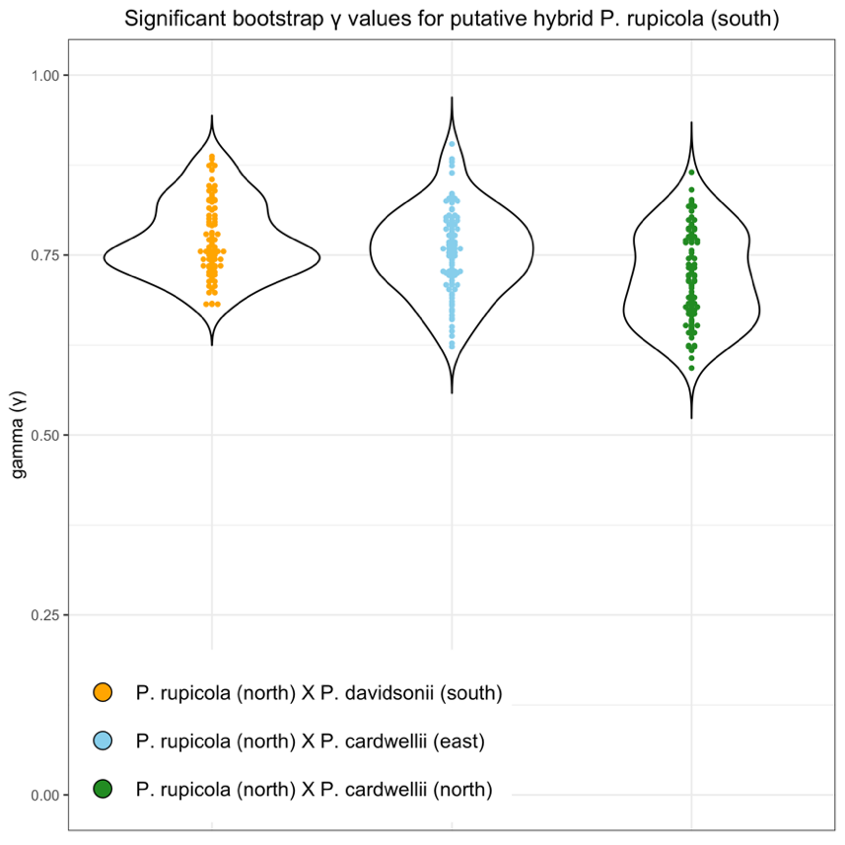

(b)

| *Supplemental Table 1.* Collection information for specimens included in the study | | | |
| --- | --- | --- | --- |
| Taxon | Latitude | Longitude | Population ID |
| *Penstemon cardwellii* | 42.32805556 | -123.8058333 | cardwellii ADW 567 |
| *Penstemon cardwellii* | 45.3027232 | -121.8184463 | cardwellii ADW 607 |
| *Penstemon cardwellii* | 44.345946 | -121.993696 | cardwellii ADW 612 |
| *Penstemon cardwellii* | 45.532477 | -122.08786 | cardwellii ADW 602 |
| *Penstemon cardwellii* | 45.5383113 | -122.097512 | cardwellii SLD 20 |
| *Penstemon cardwellii* | 42.0197251 | -123.5241447 | cardwellii SLD 11 |
| *Penstemon cardwellii* | 44.46833333 | -121.8816667 | cardwellii ADW 615 |
| *Penstemon cardwellii* | 46.314 | -122.037 | cardwellii SLD 111 |
| *Penstemon cardwellii* | 44.819678 | -122.088399 | cardwellii_BWS_108-1-4 |
| *Penstemon cardwellii* | 44.808767 | -122.110374 | cardwellii_BWS_108-5-10 |
| *Penstemon cardwellii* | 44.401918 | -121.860045 | cardwellii_BWS_109 |
| *Penstemon cardwellii* | 44.024151 | -121.804755 | cardwellii_BWS_110 |
| *Penstemon cardwellii* | 43.58548 | -122.663978 | cardwellii_BWS_112 |
| *Penstemon cardwellii* | 46.313721 | -122.036324 | cardwellii_BWS_28 |
| *Penstemon cardwellii* | 42.626077 | -123.835088 | cardwellii_BWS_90 |
| *Penstemon cardwellii* | 44.274483 | -123.579297 | cardwellii_BWS_91 |
| *Penstemon cardwellii* | 45.210899 | -123.758863 | cardwellii_BWS_92 |
| *Penstemon cardwellii* | 45.417722 | -121.825086 | cardwellii_BWS_96 |
| *Penstemon cardwellii* | 46.29557 | -122.352124 | cardwellii_BWS_97 |
| *Penstemon davidsonii* var. *davidsonii* | 37.188755 | -118.591939 | davidsonii Wilson 3554 |
| *Penstemon davidsonii* var. *davidsonii* | 44.1875 | -121.88055 | davidsonii SLD 39 |
| *Penstemon davidsonii* var. *davidsonii* | 42.944241 | -122.176547 | davidsonii_BWS_115 |
| *Penstemon davidsonii* var. *davidsonii* | 42.895602 | -122.092252 | davidsonii_BWS_116 |
| *Penstemon davidsonii* var. *davidsonii* | 43.684535 | -121.265441 | davidsonii_BWS_117 |
| *Penstemon davidsonii* var. *davidsonii* | 44.266403 | -121.787933 | davidsonii_BWS_23 |
| *Penstemon davidsonii* var. *davidsonii* | 44.021183 | -121.80011 | davidsonii_BWS_25 |
| *Penstemon davidsonii* var. *davidsonii* | 44.023102 | -121.775986 | davidsonii_BWS_27 |
| *Penstemon davidsonii* var. *davidsonii* | 42.08102 | -122.718583 | davidsonii_BWS_87 |
| *Penstemon davidsonii* var. *davidsonii* | 45.394993 | -121.658014 | davidsonii_BWS_95 |
| *Penstemon davidsonii* var. *davidsonii* | 44.0917 | -121.6467 | davidsonii_OSTL_1 |
| *Penstemon davidsonii* var. *davidsonii* | 44.4861 | -121.8386 | davidsonii_OSTL_2 |
| *Penstemon davidsonii* var. *davidsonii* | 48.832448 | -121.650946 | davidsonii_OSTL_4 |
| *Penstemon davidsonii* var. *davidsonii* | 48.734817 | -120.675033 | davidsonii_UWTL_1 |
| *Penstemon davidsonii* var. *davidsonii* | 47.444083 | -120.94285 | davidsonii_UWTL_10 |
| *Penstemon davidsonii* var. *davidsonii* | 42.555233 | -120.791667 | davidsonii_UWTL_11 |
| *Penstemon davidsonii* var. *davidsonii* | 44.0052 | -121.6424 | davidsonii_UWTL_3 |
| *Penstemon davidsonii* var. *davidsonii* | 43.68781 | -121.25558 | davidsonii_UWTL_4 |
| *Penstemon davidsonii* var. *menziesii* | 48.716888 | -121.078214 | menziesii_BWS_103 |
| *Penstemon davidsonii* var. *menziesii* | 48.706497 | -121.13966 | menziesii_BWS_104 |
| *Penstemon davidsonii* var. *menziesii* | 47.50596 | -123.2394 | menziesii_UWTL_2 |
| *Penstemon davidsonii* var. *menziesii* | 48.52361 | -120.81444 | menziesii_UWTL_5 |
| *Penstemon davidsonii* var. *menziesii* | 48.9738 | -120.19608 | menziesii_UWTL_6 |
| *Penstemon davidsonii* var. *menziesii* | 48.9363 | -120.20861 | menziesii_UWTL_7 |

*Supplemental Table 1 cont.*

| *Penstemon davidsonii* var. *menziesii* | 48.902517 | -121.466933 | menziesii_UWTL_8 |
| --- | --- | --- | --- |
| *Penstemon fruticosus* var. *fruticosus* | 47.411009 | -121.092183 | fruticosus_BWS_102 |
| *Penstemon fruticosus* var. *fruticosus* | 46.768161 | -121.67451 | fruticosus_BWS_106-1-5 |
| *Penstemon fruticosus* var. *fruticosus* | 46.906403 | -121.578231 | fruticosus_107 |
| *Penstemon fruticosus* var. *fruticosus* | 45.42266 | -121.614503 | fruticosus_BWS_93 |
| *Penstemon fruticosus* var. *fruticosus* | 44.0254 | -121.7957 | fruticosus_OSTL_10 |
| *Penstemon fruticosus* var. *fruticosus* | 44.509 | -120.6309 | fruticosus_OSTL_15 |
| *Penstemon fruticosus* var. *fruticosus* | 44.812 | -121.7934 | fruticosus_OSTL_8 |
| *Penstemon montanus* var. *montanus* | 44.511154 | -114.374046 | montanus_BWS_61-1-10 |
| *Penstemon newberryi* var. *berryi* | 40.79720833 | -123.0044444 | berryi ADW 1502 |
| *Penstemon newberryi* var. *berryi* | 40.585266 | -123.542329 | berryi_BWS_83 |
| *Penstemon newberryi* var. *berryi* | 41.389587 | -122.993862 | berryi_BWS_84-1-10 |
| *Penstemon newberryi* var. *newberryi* | 39.282751 | -120.141526 | newberryi SLD 77 |
| *Penstemon newberryi* var. *newberryi* | 38.55594 | -120.349923 | newberryi SLD 73 |
| *Penstemon newberryi* var. *newberryi* | 38.748449 | -120.051575 | newberryi SLD 69 |
| *Penstemon newberryi* var. *newberryi* | 38.197559 | -120.021536 | newberryi ADW 455 |
| *Penstemon newberryi* var. *newberryi* | 37.181157 | -118.559254 | newberryi_BWS_70 |
| *Penstemon newberryi* var. *newberryi* | 37.587992 | -118.982346 | newberryi_BWS_71 |
| *Penstemon newberryi* var. *newberryi* | 38.021527 | -119.26708 | newberryi_BWS_72 |
| *Penstemon newberryi* var. *newberryi* | 38.77672 | -119.888744 | newberryi_BWS_73 |
| *Penstemon newberryi* var. *newberryi* | 38.615967 | -119.915853 | newberryi_BWS_74 |
| *Penstemon newberryi* var. *newberryi* | 38.661708 | -120.130671 | newberryi_BWS_75 |
| *Penstemon newberryi* var. *newberryi* | 38.97312 | -120.094277 | newberryi_BWS_76 |
| *Penstemon newberryi* var. *newberryi* | 39.947764 | -121.140946 | newberryi_BWS_79 |
| *Penstemon newberryi* var. *newberryi* | 39.880031 | -121.161161 | newberryi_BWS_80 |
| *Penstemon newberryi* var. *newberryi* | 41.231111 | -122.382305 | newberryi_BWS_85 |
| *Penstemon rupicola* | 42.48666667 | -124.02 | rupicola ADW 575 |
| *Penstemon rupicola* | 45.5383113 | -122.097512 | rupicola SLD 18 |
| *Penstemon rupicola* | 42.234281 | -123.791754 | rupicola SLD 14 |
| *Penstemon rupicola* | 47.268312 | -121.366167 | rupicola_BWS_100 |
| *Penstemon rupicola* | 47.411009 | -121.092183 | rupicola_BWS_101 |
| *Penstemon rupicola* | 46.778033 | -121.760309 | rupicola_BWS_105 |
| *Penstemon rupicola* | 46.768161 | -121.67451 | rupicola_BWS_106-6 |
| *Penstemon rupicola* | 44.024151 | -121.804755 | rupicola_BWS_111 |
| *Penstemon rupicola* | 43.09002 | -122.249755 | rupicola_BWS_113 |
| *Penstemon rupicola* | 40.658657 | -123.219845 | rupicola_BWS_81 |
| *Penstemon rupicola* | 41.223113 | -122.37963 | rupicola_BWS_86 |
| *Penstemon rupicola* | 42.236884 | -123.797584 | rupicola_BWS_89 |
| *Penstemon rupicola* | 47.436029 | -121.778519 | rupicola_BWS_98 |
| *Penstemon rupicola* | 47.333277 | -121.384662 | rupicola_BWS_99 |
| *Penstemon rupicola* | 44.0254 | -121.7957 | rupicola_OSTL_5 |
| *Penstemon rupicola* | 44.6475 | -122.0628 | rupicola_OSTL_6 |
| *Penstemon rupicola* | 42.8036 | -122.2605 | rupicola_OSTL_7 |
| *Penstemon rupicola* | 47.4388 | -121.7726 | rupicola_UWTL_23 |

| *Supplemental Table 2.* Average variable importance for each model included in ensemble for SDMs | | | | | | | |
| --- | --- | --- | --- | --- | --- | --- | --- |
| *P. rupicola* | bio_4 | bio_5 | bio_8 | bio_14 | bio_15 | bio_18 | bio_19 |
| MAXENT.Phillips | 0.143 | 0.197 | 0.532 | 0.289 | 0.206 | 0.217 | 0.809 |
| GLM | 0.449 | 0.069 | 0.449 | 0.452 | 0.152 | 0.063 | 0.883 |
| RF | 0.055 | 0.035 | 0.220 | 0.151 | 0.102 | 0.035 | 0.417 |
| GBM | 0.006 | 0.012 | 0.572 | 0.051 | 0.460 | 0.003 | 0.747 |
| *P. newberryi* | bio_4 | bio_5 | bio_8 | bio_14 | bio_15 | bio_18 | bio_19 |
| MAXENT.Phillips | 0.178 | 0.303 | 0.244 | 0.116 | 0.264 | 0.568 | 0.211 |
| GLM | 0.355 | 0.295 | 0.200 | 0.069 | 0.179 | 0.656 | 0.151 |
| RF | 0.380 | 0.056 | 0.400 | 0.005 | 0.225 | 0.323 | 0.028 |
| GBM | 0.569 | 0.002 | 0.292 | 0.000 | 0.402 | 0.399 | 0.002 |
| *P. fruticosus* | bio_4 | bio_5 | bio_8 | bio_14 | bio_15 | bio_18 | bio_19 |
| MAXENT.Phillips | 0.276 | 0.046 | 0.714 | 0.352 | 0.294 | 0.138 | 0.187 |
| GLM | 0.523 | 0.027 | 0.112 | 0.449 | 0.206 | 0.428 | 0.029 |
| RF | 0.190 | 0.088 | 0.538 | 0.222 | 0.248 | 0.272 | 0.185 |
| GBM | 0.320 | 0.009 | 0.548 | 0.080 | 0.038 | 0.074 | 0.203 |
| *P. davidsonii* | bio_4 | bio_5 | bio_8 | bio_14 | bio_15 | bio_18 | bio_19 |
| MAXENT.Phillips | 0.324 | 0.203 | 0.329 | 0.136 | 0.157 | 0.179 | 0.073 |
| GLM | 0.341 | 0.311 | 0.038 | 0.110 | 0.102 | 0.544 | 0.381 |
| RF | 0.107 | 0.050 | 0.105 | 0.029 | 0.087 | 0.044 | 0.055 |
| GBM | 0.519 | 0.058 | 0.428 | 0.017 | 0.094 | 0.011 | 0.010 |
| *P. cardwellii* | bio_4 | bio_5 | bio_8 | bio_14 | bio_15 | bio_18 | bio_19 |
| MAXENT.Phillips | 0.080 | 0.005 | 0.207 | 0.047 | 0.269 | 0.088 | 0.739 |
| GLM | 0.801 | 0.379 | 0.739 | 0.364 | 0.296 | 0.230 | 0.649 |
| RF | 0.043 | 0.043 | 0.090 | 0.201 | 0.167 | 0.048 | 0.252 |
| GBM | 0.006 | 0.005 | 0.337 | 0.080 | 0.465 | 0.005 | 0.795 |
| Models correspond to the Maximum Entropy model as implemented in Maxent (MAXENT.Phillips), General Linear Models (GLM), Random Forests (RF), and Generalized Boosting Models (GBM). Variable names correspond to bioclim variables: bio4 = Temperate Seasonality (standard deviation x 100); bio5 = Max Temperature of Warmest Month; bio8 = Mean Temperature of Wettest Quarter; bio14 = Precipitation of Driest Month; bio15 = Precipitation Seasonality (Coefficient of Variation); bio18 = Precipitation of Warmest Quarter; bio19 = Precitipitation of Coldest Quarter. | | | | | | | |

| *Supplemental Table 3.* Error rates and Confusion Matrix for each replicate *delimitR* run for *P. rupicola* | | | | | | | | | | | | | |
| --- | --- | --- | --- | --- | --- | --- | --- | --- | --- | --- | --- | --- | --- |
| Error: | OOB | Model 1 | Model 2 | Model 3 | Model 4 | Model 5 | Model 6 | Model 7 | Model 8 | Model 9 | Model 10 | Model 11 | Model 12 |
| rep 0 | 0.194 | 0.077 | 0.087 | 0.040 | 0.360 | 0.329 | 0.359 | 0.050 | 0.157 | 0.094 | 0.241 | 0.286 | 0.250 |
| rep 1 | 0.193 | 0.068 | 0.088 | 0.043 | 0.362 | 0.324 | 0.358 | 0.049 | 0.159 | 0.096 | 0.238 | 0.285 | 0.245 |
| rep 2 | 0.192 | 0.072 | 0.085 | 0.044 | 0.361 | 0.322 | 0.353 | 0.049 | 0.153 | 0.094 | 0.241 | 0.284 | 0.242 |
| rep 3 | 0.192 | 0.073 | 0.087 | 0.040 | 0.355 | 0.324 | 0.354 | 0.049 | 0.156 | 0.093 | 0.242 | 0.281 | 0.250 |
| rep 4 | 0.192 | 0.075 | 0.085 | 0.040 | 0.356 | 0.320 | 0.353 | 0.051 | 0.158 | 0.097 | 0.240 | 0.281 | 0.245 |
| Votes: | PP | Model 1 | Model 2 | Model 3 | Model 4 | Model 5 | Model 6 | Model 7 | Model 8 | Model 9 | Model 10 | Model 11 | Model 12 |
| rep 0 | 0.724 | 36 | 0 | 0 | 79 | 338 | 47 | 0 | 0 | 0 | 0 | 0 | 0 |
| rep 1 | 0.669 | 87 | 0 | 0 | 92 | 250 | 71 | 0 | 0 | 0 | 0 | 0 | 0 |
| rep 2 | 0.761 | 15 | 0 | 0 | 89 | 370 | 26 | 0 | 0 | 0 | 0 | 0 | 0 |
| rep 3 | 0.424 | 48 | 0 | 0 | 69 | 299 | 84 | 0 | 0 | 0 | 0 | 0 | 0 |
| rep 4 | 0.475 | 100 | 0 | 0 | 92 | 116 | 192 | 0 | 0 | 0 | 0 | 0 | 0 |
| The best model for each replicate is shaded in grey. Model votes are out of 500 *fastsimcoal* replicates. Each model includes three populations. OOB = out-of-bag error rate; PP = posterior probability. Models receiving at least 5% of votes in at least one replicate are as follows: model 1 = post-LGM divergence and north/south populations sister; model 4 = pre-LGM divergence, north/south sister, and no migration; model 5 = pre-LGM divergence, north/south sister, and migration between the Klamath and southern populations; model 6 = pre-LGM divergence, north/south sister, and migration between the Klamath and north populations. | | | | | | | | | | | | | |

| *Supplemental Table 4.* Error rates and Confusion Matrix for each replicate *delimitR* run for *P. cardwellii* | | | | | | | | | | | | | |
| --- | --- | --- | --- | --- | --- | --- | --- | --- | --- | --- | --- | --- | --- |
| Error: | OOB | Model 1 | Model 2 | Model 3 | Model 4 | Model 5 | Model 6 | Model 7 | Model 8 | Model 9 | Model 10 | Model 11 | Model 12 |
| rep 0 | 0.043 | 0.009 | 0.009 | 0.008 | 0.012 | 0.058 | 0.089 | 0.007 | 0.054 | 0.057 | 0.022 | 0.095 | 0.099 |
| rep 1 | 0.043 | 0.009 | 0.009 | 0.009 | 0.012 | 0.055 | 0.088 | 0.008 | 0.054 | 0.054 | 0.024 | 0.097 | 0.102 |
| rep 2 | 0.043 | 0.010 | 0.010 | 0.008 | 0.013 | 0.055 | 0.089 | 0.007 | 0.051 | 0.055 | 0.023 | 0.095 | 0.101 |
| rep 3 | 0.044 | 0.010 | 0.011 | 0.009 | 0.013 | 0.056 | 0.090 | 0.007 | 0.055 | 0.056 | 0.024 | 0.091 | 0.105 |
| rep 4 | 0.043 | 0.010 | 0.009 | 0.007 | 0.013 | 0.058 | 0.088 | 0.006 | 0.053 | 0.054 | 0.022 | 0.094 | 0.098 |
| Votes: | PP | Model 1 | Model 2 | Model 3 | Model 4 | Model 5 | Model 6 | Model 7 | Model 8 | Model 9 | Model 10 | Model 11 | Model 12 |
| rep 0 | 0.985 | 122 | 329 | 39 | 0 | 2 | 1 | 0 | 5 | 0 | 0 | 0 | 2 |
| rep 1 | 0.991 | 114 | 351 | 27 | 0 | 1 | 3 | 1 | 1 | 1 | 0 | 1 | 0 |
| rep 2 | 0.998 | 122 | 352 | 16 | 0 | 3 | 2 | 0 | 1 | 2 | 0 | 1 | 1 |
| rep 3 | 0.982 | 93 | 369 | 27 | 0 | 1 | 3 | 0 | 2 | 3 | 0 | 0 | 2 |
| rep 4 | 0.979 | 180 | 263 | 48 | 1 | 2 | 1 | 1 | 1 | 1 | 0 | 0 | 2 |
| The best model for each replicate is shaded in grey. Model votes are out of 500 *fastsimcoal* replicates. Each model includes three populations. OOB = out-of-bag error rate; PP = posterior probability. Models receiving at least 5% of votes in at least one replicate are as follows: model 1 = post-LGM divergence and Klamath/east populations sister; model 2 = post-LGM divergence and west/east sister; model 3 = post-LGM divergence and Klamath/west sister. | | | | | | | | | | | | | |

| *Supplemental Table 5.* Error rates and Confusion Matrix for each replicate *delimitR* run for *P. newberryi* | | | | |
| --- | --- | --- | --- | --- |
| Error: | OOB | Model 1 | Model 2 | Model 3 |
| rep 0 | 0.049 | 0.063 | 0.003 | 0.082 |
| rep 1 | 0.047 | 0.061 | 0.002 | 0.077 |
| rep 2 | 0.048 | 0.062 | 0.002 | 0.079 |
| rep 3 | 0.047 | 0.063 | 0.003 | 0.075 |
| rep 4 | 0.046 | 0.062 | 0.002 | 0.075 |
| Votes: | PP | Model 1 | Model 2 | Model 3 |
| rep 0 | 0.957 | 453 | 0 | 47 |
| rep 1 | 0.962 | 459 | 0 | 41 |
| rep 2 | 0.989 | 478 | 0 | 22 |
| rep 3 | 0.986 | 460 | 0 | 40 |
| rep 4 | 0.800 | 382 | 0 | 118 |
| The best model for each replicate is shaded in grey. Model votes are out of 500 *fastsimcoal* replicates. Each model includes two populations. OOB = out-of-bag error rate; PP = posterior probability. Models receiving at least 5% of votes in at least one replicate are as follows: model 1 = post-LGM divergence; model 3 = pre-LGM divergence with migration. | | | | |

| *Supplemental Table 6.* Error rates and Confusion Matrix for each replicate *delimitR* run for *P. davidsonii* | | | | |
| --- | --- | --- | --- | --- |
| Error: | OOB | Model 1 | Model 2 | Model 3 |
| rep 0 | 0.062 | 0.118 | 0.002 | 0.065 |
| rep 1 | 0.058 | 0.111 | 0.002 | 0.060 |
| rep 2 | 0.060 | 0.116 | 0.002 | 0.062 |
| rep 3 | 0.059 | 0.114 | 0.002 | 0.062 |
| rep 4 | 0.058 | 0.114 | 0.002 | 0.060 |
| Votes: | PP | Model 1 | Model 2 | Model 3 |
| rep 0 | 0.997 | 444 | 2 | 54 |
| rep 1 | 0.995 | 459 | 0 | 41 |
| rep 2 | 0.994 | 404 | 0 | 96 |
| rep 3 | 0.994 | 372 | 0 | 128 |
| rep 4 | 0.993 | 408 | 1 | 91 |
| The best model for each replicate is shaded in grey. Model votes are out of 500 *fastsimcoal* replicates. Each model includes two populations. OOB = out-of-bag error rate; PP = posterior probability. Models receiving at least 5% of votes in at least one replicate are as follows: model 1 = post-LGM divergence; model 3 = pre-LGM divergence with migration. | | | | |

| *Supplemental Table 7.* Error rates and Confusion Matrix for each replicate *delimitR* run for *P. fruticosus* | | | | |
| --- | --- | --- | --- | --- |
| Error: | OOB | Model 1 | Model 2 | Model 3 |
| rep 0 | 0.200 | 0.302 | 0.014 | 0.283 |
| rep 1 | 0.207 | 0.315 | 0.014 | 0.291 |
| rep 2 | 0.202 | 0.307 | 0.014 | 0.287 |
| rep 3 | 0.205 | 0.313 | 0.015 | 0.287 |
| rep 4 | 0.198 | 0.305 | 0.015 | 0.274 |
| Votes: | PP | Model 1 | Model 2 | Model 3 |
| rep 0 | 0.466 | 308 | 1 | 191 |
| rep 1 | 0.799 | 451 | 6 | 43 |
| rep 2 | 0.094 | 113 | 338 | 49 |
| rep 3 | 0.727 | 70 | 2 | 428 |
| rep 4 | 0.174 | 150 | 6 | 344 |
| The best model for each replicate is shaded in grey. Model votes are out of 500 *fastsimcoal* replicates. Each model includes two populations. OOB = out-of-bag error rate; PP = posterior probability. Models receiving at least 5% of votes in at least one replicate are as follows: model 1 = post-LGM divergence; model 2 = pre-LGM divergence and no migration; model 3 = pre-LGM divergence with migration. | | | | |

| *Supplemental Table 8.* *HyDe* summary statistics | | | |
| --- | --- | --- | --- |
|  | Tests considered | | Significant results |
| Total | 423 | 45 (10.6%) | |
| Tests including Klamath lineages | 273 (64.5%) | 39 (86.7%) | |
|  | Significant results (individual tests) | | |
| Tests including southern *P. davidsonii* | 82/100 (82.0%) | | |
| Tests including southern *P. rupicola* | 30/44 (68.2%) | | |
